# Distribution-driven disturbance of a molecular community: a theoretical framework

**DOI:** 10.1101/2024.03.11.584481

**Authors:** Gustavo A. Ramírez, Gabriel C. Mel

## Abstract

An important and understudied area in microbial ecology concerns the very large number of very rare species within an environment. Study of such species based on non-specific sequencing of environmental samples is practically difficult simply due to their rarity. A previous report demonstrated that pretreating samples with the photoactive dye PMA before subsequent PCR and sequencing can increase sensitivity to rare species. Here we propose a mathematical model based on equilibrium chemical thermodynamics that explains this effect and suggests a simple quantitative experiment.

## Motivation and approach

In nature, microbial communities are comprised of distinct lineages (ie. species) of unequal abundance. Determining microbial community diversity and structure is of central importance to understanding the ecological function and evolutionary underpinnings of a habitat. Often, only a few lineages represent the majority of microorganisms in a given community. Despite this, thousands of distinct lineages may co-exist in any given environment, meaning that rare lineages, those with low cellular abundances, represent the most diverse part of a natural community.

Assuming that the DNA extract pool from a given environmental sample is an accurate representation of the source microbial community, a skewed molecular distribution, where most of the DNA in the extract pool represents a few dominant microbial lineages, can be expected. Thus, our understanding of any given environment through next-generation (NGS) DNA sequencing approaches will be naturally skewed toward the most numerically dominant lineages. Rare lineages with unexplored and potentially important ecological functions are generally neglected by this widespread technology.

Here, we generate a theoretical framework for exploring molecular information from rare microbial lineages using a modified Polymerase Chain Reaction (PCR) protocol. Specifically, we model the effects of DNA “silencing” by Propidium Monoazide (PMA), a photoactive dye, which indiscriminately binds DNA and, when activated by light, covalently bonds the double helix, thereby preventing PCR-based amplification and ultimately downstream sequencing [1]. We posit, based on a chemical equilibrium model, that non-targeted silencing of a total DNA pool extracted from a natural microbial community will result in a higher rate of silencing of DNA representing numerically dominant lineages, consistent with the findings of [2]. Below, we detail this model-based prediction, which in turn suggests a simple experimental approach for empirical validation.

## A simple equilibrium model

Assume a solution with several DNA sequences representing a total DNA extraction from an environmental microbial community, and consider the problem of amplifying the *rarest* species. With ordinary PCR and minimal to no primer bias [3], one expects to roughly double the number of target templates each cycle, leading to amplification to roughly 2, 4, 8, … template copies as cycles are performed, or more generally a factor of 2*n* for *n* cycles. Ideally, using universal 16S rRNA gene primers, amplification should not depend on the identity of the DNA sequence. Thus, we expect that all sequences experience roughly the same amplification factor, and so the relative concentrations of any two sequences is unchanged:

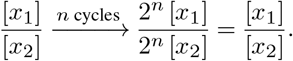

That is, PCR amplification of all primed gene targets in the bulk DNA template should not directly affect the profile of sequences present. Rare target sequences, assuming they are flanked by the primer probes, are no more and no less rare (relative to the full population) after *n*-cycles of PCR. Under these assumptions, i) the primary determinant for the relative concentrations of any two sequence types amplified with equal efficiency is their relative prevalence at the onset of PCR, and ii) if extremely abundant targets are “eliminated” at onset, rare sequences will increase in relative abundance. By fusing double-stranded DNA, PMA treatment allows one to effectively remove, or silence, hybridized sequences from the initial DNA template (Figure 1). Specifically, PMA intercalates double-stranded DNA. When the DNA-PMA sample is irradiated with visible light, PMA-photo activation promotes the irreversible formation of a covalent bond between the strands of the DNA double helix [1], inhibiting DNA denaturation during PCR and ultimately disrupting amplification of the corresponding gene target. This gives a method to selectively target common sequences since, due to their relatively high prevalence, these will generally be more hybridized, and thus more vulnerable to PMA intercalation and eventual PCR silencing.

**Fig. 1.**
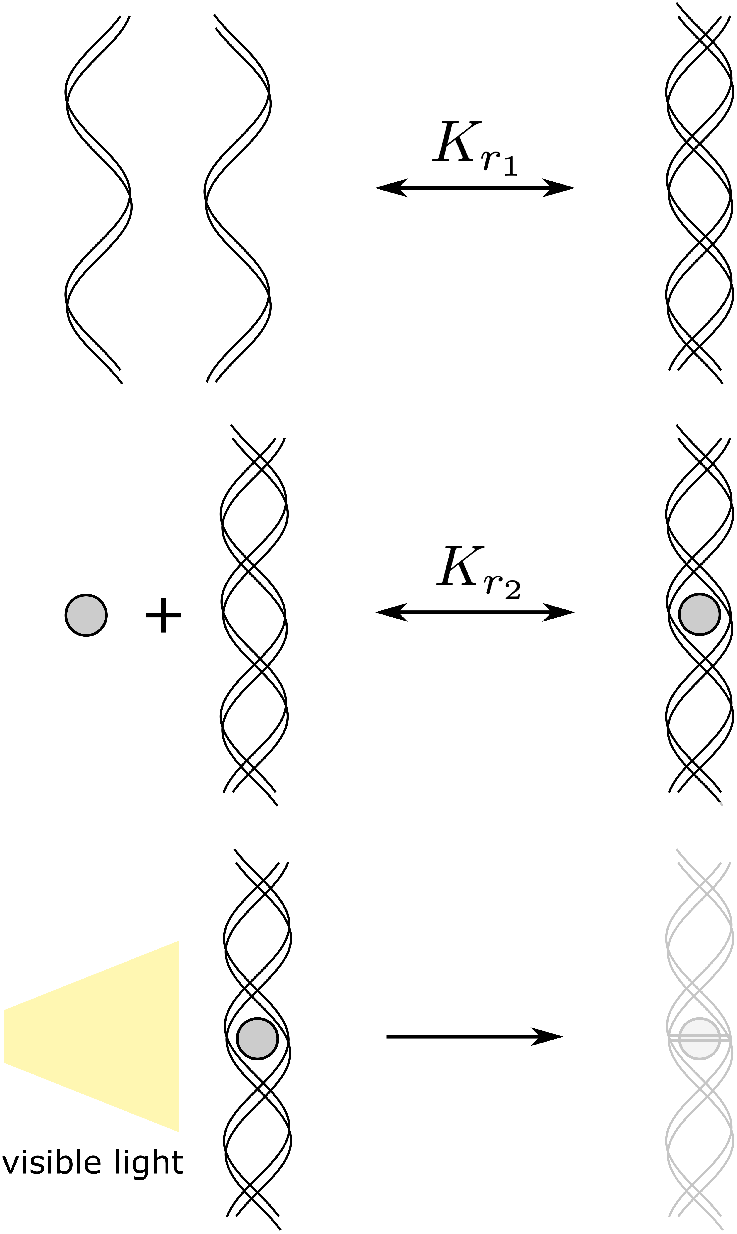
Illustration of PMA treatment. PMA intercalates hybridized sequences (bottom left), and when irradiated, covalently fuses the two strands, rendering the sequences inert with respect to PCR (right).

To see this, choose a single DNA target sequence (eg. a 16S rRNA gene fragment) and let [*x*], [*x*_2_] represent its unhybridized (single stranded *(ss)*DNA) and hybridized (double stranded *(ds)*DNA) concentrations, respectively. Denote by *µ* := 2 [*x*_2_] + [*x*] the total count of the target sequence (in a solution of fixed volume, species counts and concentrations are proportional, and we will use the terms interchangeably). Further, let [*P*] be the concentration of PMA, and [*x*_2*P*_] be the concentration of hybridized targets with PMA intercalated. At the moment of irradiation, let us assume the solution is at equilibrium for both of the reactions

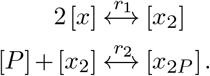

Equilibrium implies

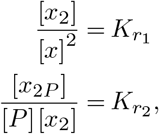

where 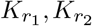 are the temperature-dependent equilibrium constants of the two reactions (Figure 1). After irradiation, intercalated strands are irreversibly fused by covalent bonds and are rendered inert, or silenced, with respect to PCR, so the effective concentration *µ*_*eff*_ of the sequence becomes

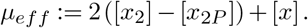

Let us consider the effect of PMA in terms of the reduction coefficient,

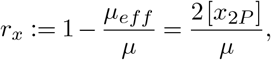

which corresponds to the fraction of total sequence count that will be eliminated by PMA treatment.

We can express the total concentration *µ* at the moment before irradiation as *µ* = 2 [*x*_2_] + 2 [*x*_2*P*_] + [*x*]. Thus, together with the equilibrium conditions, we have the following system of equations relating [*x*], [*x*_2_], and [*x*_2*P*_]:

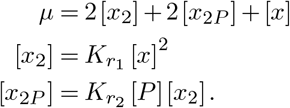

Eliminating [*x*], [*x*_2*P*_] and solving, we find

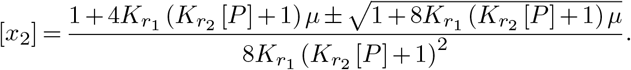

Since 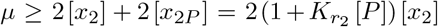, we find that only the negative solution is possible. Substituting into 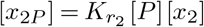 yields [*x*_2*P*_] and using 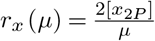, we obtain

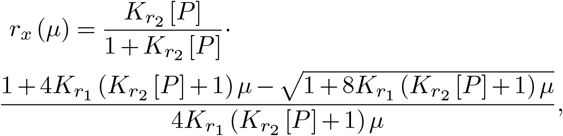

which gives the reduction factor *r*_*x*_ (*µ*) associated with a species of concentration *µ*. Let us make some simple observations about this function. First, *r*_*x*_ is a monotonically increasing function of *µ*, which means common species are removed more than rare ones. As *µ* → 0 (ie. for rare species), *r*_*x*_ → 0, indicating that rare species are essentially unaffected by the process. As *µ* → ∞ (ie. for common species), 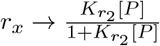, which approaches 1 (ie. total removal) as the concentration of PMA is made large. Defining

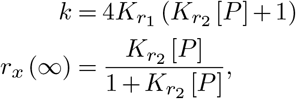

we can write the reduction as

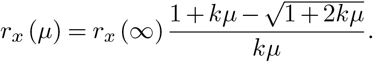

Figure 2 plots this function *r*_*x*_ (*µ*) on a logarithmic axis. More common species experience a greater reduction factor. The parameters *k, r*_*x*_ (∞) (which in turn depend on the equilibrium constants and concentration of PMA) have a simple effect on the curve: *r*_*x*_ (∞) simply scales the curve vertically, while *k* shifts the curve horizontally, moving the threshold where the reduction factor transitions low to high. For an interactive demonstration of the behavior of *r*_*x*_ as a function of the species concentration *µ* and the parameters *k, r*_*x*_(∞), see this figure created using the Desmos online graphing software.

**Fig. 2.**
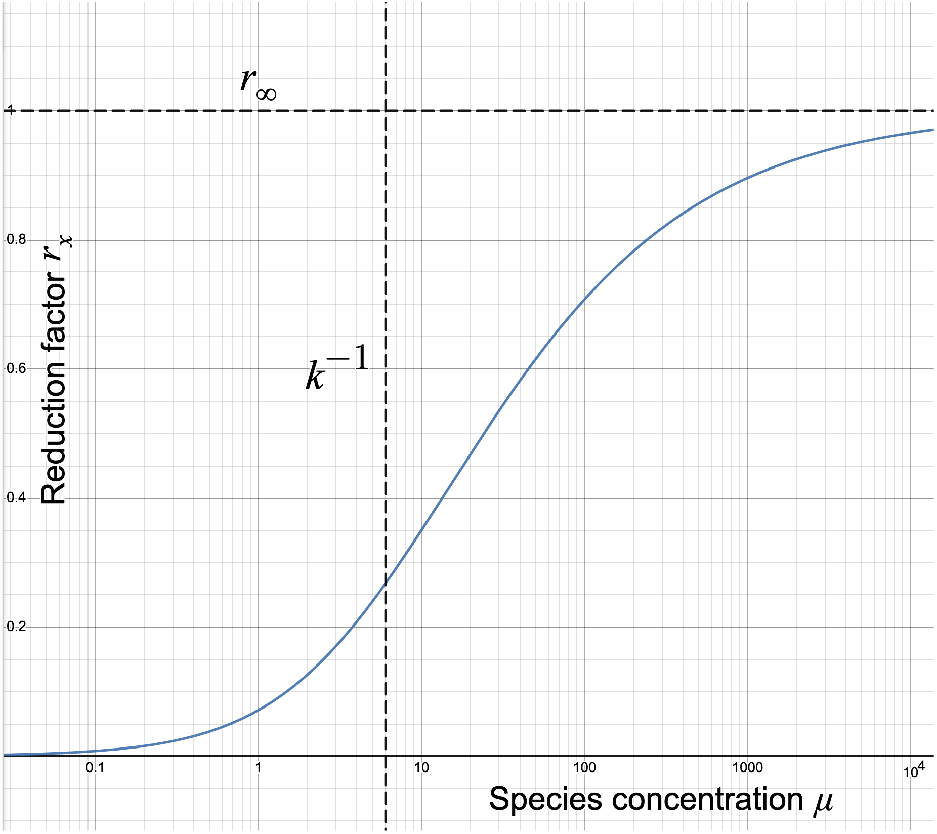
Graph of reduction ratio *r*_*x*_ as a function of species concentration (or count) *µ*. Common species are reduced more than rare ones. The reduction coefficient transitions from low to high (with an asymptotic value of *r*_*x*_(∞), here chosen to be 1) around the threshold concentration *k*^−1^.

## Discussion

In this work we propose a simple explanation for the rare species amplification effect of PMA treatment previously observed in [2]. Our goal is not to produce a comprehensive biochemical model capturing all details of PMA and PCR chemistry, but rather to illustrate how simple, well understood chemical thermodynamic mechanisms can account for the observed effect. Quantitative PCR experiments involving multiple target sequences with controlled concentrations should allow one to test the predictions of the model, in particular the reduction coefficient *r*_*x*_(*µ*). Within a reasonable range of variability, target-dependent sequence lengths, hybridization equilibrium constants, or primer affinities would affect quantitative predictions, but would leave the qualitative behavior relatively unchanged. We leave these for future work.

